# AI Assisted Native Proteomics: Delineating Ribosomal Protein Conformations Pre- and Post-Assembly

**DOI:** 10.1101/2024.11.09.622757

**Authors:** Wenjing Zhang, Chen Sun, Zhang Xu, Wei Xu

## Abstract

The simultaneous identification of proteins as well as their conformations in a biological system would greatly enhance our understanding of cellular mechanisms and disease. As an emerging technique, native proteomics analyzes proteins in their native states, facilitating the acquisition of protein stoichiometry, post-translational modifications (PTMs), and interactions with ligands. However, revealing protein conformations at the proteome scale remains a significant challenge. In this study, we propose an AI-assisted native proteomics method that integrates a protein structure prediction (PSP) module with top-down proteomics (TDP) and native mass spectrometry (nMS) to acquire both proteome identity and conformations. First, protein sequences and PTMs are obtained using the TDP method, while protein solvent-accessible surface area (SASA) is measured in parallel by nMS. These data are then input into the PSP module to acquire the most probable conformation of the protein under nMS experimental conditions. We validate the feasibility and accuracy of this method through the analysis of both a globular protein and an intrinsically disordered protein. Additionally, we apply this approach to delineate the conformations of ribosomal proteins pre- and post-assembly. Interactions of ribosomal proteins with drug molecules are also explored. By enabling proteome identification and conformation characterization from relatively small amounts of endogenous proteins, this method would bridge the gap between structural biology and conventional proteomics technologies.

## Introduction

Proteins are the primary functional molecules in cells, responsible for most cellular processes.^1,2^ Mass spectrometry (MS) based bottom-up and top-down proteomics techniques have enabled large-scale study of proteomes expressed by a genome.^3-7^ With the ability to acquire protein sequences and post translational modifications (PTM) in a biological system, proteomics is a powerful tool in biology that enhances our understanding of cellular functions, disease mechanisms,^8^ and drug development.^7^ However, protein functions also heavily depend on their structures, and protein misfolding is a prevalent factor linked to a diverse array of pathologies.^9-11^ Conventional technologies for protein structural determination such as X-ray crystallography, NMR spectroscopy and cryo-electron microscopy are high-resolution and low-throughput techniques that require large amounts of purified and homogeneous samples.^12-14^

Facing these challenges, many techniques have been developed and coupled with MS to probe protein structures in a high throughput fashion.^15-17^ Hydrogen-deuterium exchange MS (HDX-MS) enables the investigation of protein conformational dynamics and interactions, providing insights into protein transient states and flexible regions.^18-20^ Cross-linking mass spectrometry (XL-MS) uses chemical cross-linkers to covalently link interacting proteins or regions within a protein, effectively “freezing” their spatial relationships.^21,22^ Without the need of antibodies, proximity labeling can be conducted directly within living cells under natural conditions.^23,24^ These techniques are advantageous for capturing transient or weak protein interactions that occur in vivo, helping researchers better understand the complex biological processes within cells.^25,26^ Using softer ionization methods, native MS (nMS) analyzes proteins in their native, functional states, allowing the study of protein folding, assembly, and interactions with ligands.^27-29^ It is also found that the charge state distributions (CSDs) of proteins in a native mass spectrum correlate with their solvent accessible surface area (SASA).^30-36^ As an emerging and fast-evolving technique, native proteomics involves proteome characterization under native conditions, ultimately to acquire both protein identity (sequence and PTM) and structural information in a high-throughput fashion.^37-42^

AI-assisted protein structure prediction has revolutionized the field of structural biology by leveraging advanced computational methods to predict protein structures from amino acid sequences.^43-46^ By integrating extensive datasets from experimental structures, AI models can learn complex patterns, achieving remarkable accuracy in predicting protein folding and interactions.^47-50^ However, proteins exhibit significant flexibility and dynamics, often existing in diverse microenvironments.^51-53^ Therefore, incorporating structural information under experimental conditions into AI models can significantly enhance prediction accuracy and broaden their applications.^54-57^ For instance, integrating distance restraints from XL-MS has enabled comprehensive descriptions of intrinsically disordered proteins (IDPs).^54,58-60^

In this study, we developed an AI-assisted native proteomics method aimed at acquiring protein sequence, post-translational modification (PTM), and conformational information simultaneously in a high-throughput manner. Specifically, proteins extracted under native conditions are divided into two parts: protein identity (including sequences and PTMs) is obtained through a top-down mass spectrometry (TDMS) experiment, while conformational information (specifically SASA), is acquired using nMS. Both protein identity and conformational data are then inputted into a PSP AI module (Structure-to-Structure Translation, STR2STR),^61,62^ allowing us to determine the most probable conformations of each protein under the experimental conditions. After validating this method with model proteins, we applied it to investigate the conformations of ribosomal proteins before and after they form the ribosome complex. Results show that ribosomal proteins with intrinsically disordered regions (IDRs) typically exhibit multiple conformation ensembles in their monomer state, while adopting a single conformation upon forming the ribosome complex. Further experiments demonstrate that these multiple conformations can enhance interactions with drug molecules. Therefore, when designing drugs aimed at blocking complex formation, it is crucial to target protein conformations in their monomer state rather than those in the complex. Potentially, this method would bridge the gap between conventional proteomics and structural biology techniques.

## Results and discussions

### Workflow of PSP-native proteomics

As illustrated in Figure 1a, proteins extracted from biological samples under nondenaturing conditions are divided into two portions. The first portion undergoes RPLC-TDMS analysis (Figure 1b) to identify proteome sequences and their post-translational modifications (PTMs). This analysis employs a data-independent acquisition (DIA) method to collect high-quality tandem mass spectra by fragmenting all ions within a RPLC fraction using high-energy collision dissociation (HCD). Details of the DIA-TDMS strategy are provided in a parallel manuscript.^63^ The second portion of the protein sample is analyzed using SEC-nMS (Figure 1c). In this step, proteomes are separated and analyzed under native conditions, with protein solvent-accessible surface areas (SASAs) determined from the charge state distributions (CSDs) of protein ions in the native mass spectra. In step 4, the protein sequence obtained from step 2 is inputted into the PSP AI module, which generates a set of possible structures (conformations) for the protein. The SASA information of the same protein (obtained from step 3) is then used to constrain these conformations. The structure with a SASA that most closely matches the experimental results is selected as the most probable conformation of the protein.

**Figure 1.**
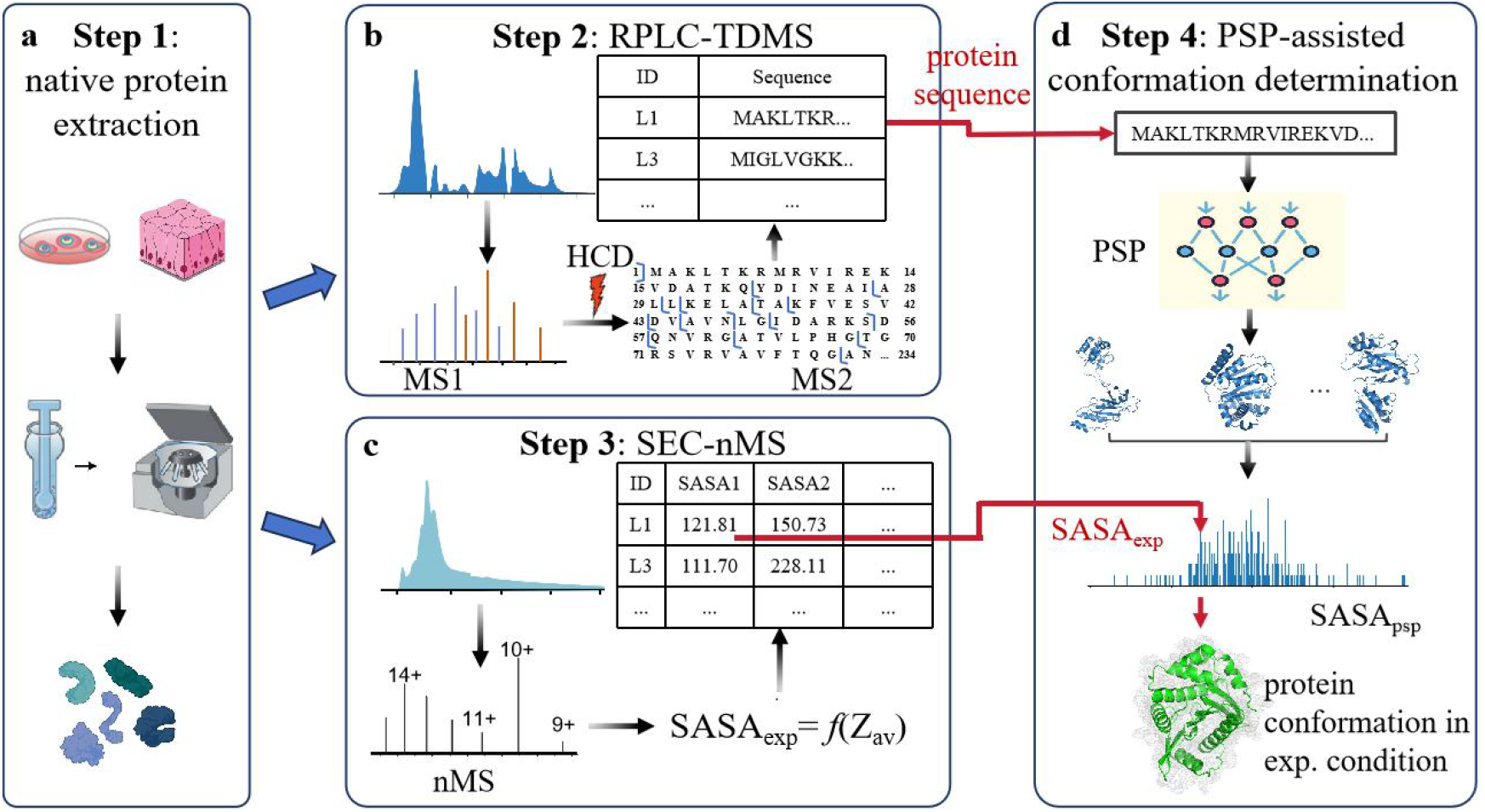
Schematic flowchart of PSP-native proteomics. (a) Protein sample extraction under nondenaturing conditions. (b) RPLC-TDMS analysis of the first sample portion. (c) SEC-nMS experiment to obtain protein SASAs. (d) PSP-assisted protein conformation determination.

### Analysis of model proteins

To validate the PSP assisted native proteomics method, we analyzed two model proteins: α-Syn, representative of intrinsically disordered proteins (IDPs), and Lys, a globular protein. Purified α-Syn and Lys were dissolved separately in 100 mM ammonium acetate buffer. Since no separation was needed for each protein sample, RPLC was not used in step 2. Figure 2a plots the tandem mass spectrum (MS/MS) of α-Syn, along with the corresponding sequence information. SEC-nMS revealed a native mass spectrum (Figure 2b) with two distinct CSDs. Unlike globular proteins, which typically adopt a single stable conformation, IDPs are highly flexible and usually exist as conformational ensembles. The observation of two distinct CSDs in the native mass spectrum suggests that α-Syn may exist as two distinct conformational ensembles under the given solvent conditions. Calculation of the SASA yielded values of 91.23 nm^2^ for the low CSD and 151.89 nm^2^ for the high CSD. Using the PSP module to predict possible conformational ensembles from the α-Syn sequence, these SASA values were then employed to identify the corresponding conformations. As depicted in Figure 2c, we obtained both a compact conformation (in red) and an open conformation (in green) for α-Syn. The open conformation aligns well with the PED00024 structure from the Protein Ensemble Database (PED),^64^ which is the well-known monomer structure tending to form oligomers and amyloid fibrils. Moreover, the compact conformation agrees with PDBDEV_00000082 structure from fluorescence resonance energy transfer (FRET) experiments.^65^ The presence of both compact and open conformations in α-Syn is consistent with previous studies of IDPs, such as those showing that calmodulin (CaM) exhibits both open and closed states, with a dynamic equilibrium between these conformations occurring on time scales ranging from milliseconds to microseconds.^66^ It is hypothesized that the lower protein concentration (for example, much lower than that in NMR conditions) may contribute to the existence of the compact conformation with a relatively high percentage. In contrast, Figures 2d-f show the results for Lys, where a single compact conformation was observed. Agreements between the conformations obtained using the proposed method and those reported in other experimental databases suggest that this PSP-assisted native proteomics method is effective for studying both IDP and globular proteins.

**Figure 2.**
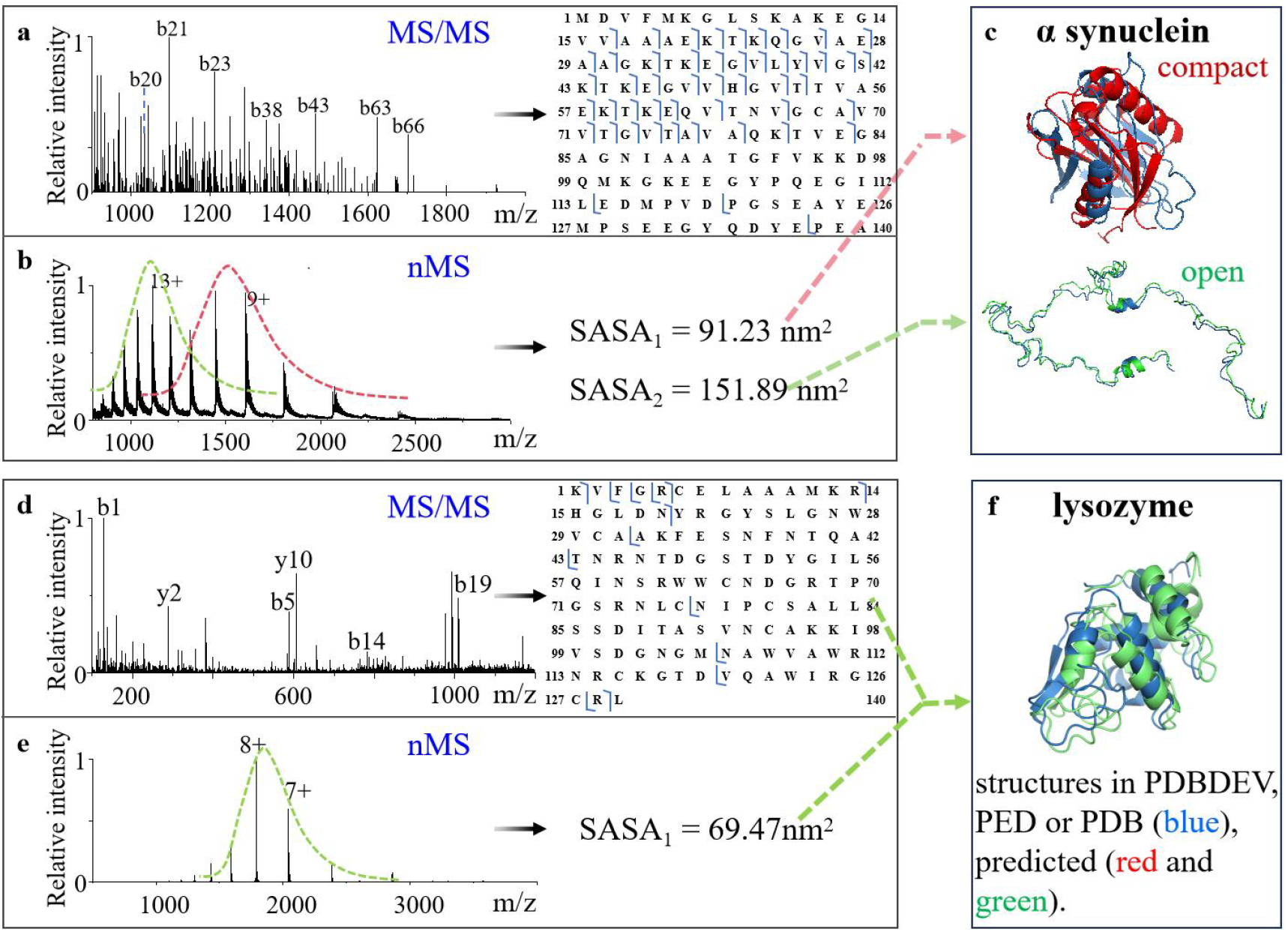
Analysis of model proteins: α-Syn (a-c) and Lys (d-f). Sequence determination of α-Syn (a) and Lys (d) by tandem MS. SASA measurement of α-Syn (b) and Lys (e) by native MS. (c) The two most probable conformations of α-Syn obtained by the proposed method (red and green) and the corresponding structures (blue) of α-Syn in PDB-Dev^67-71^ (https://pdb-dev.wwpdb.org) and PED^72^ (https://proteinensemble.org). (f) Comparison of the conformation of Lys acquired from the proposed method (green) and that in the PDB database (PDB ID: 7CDK, blue).

### Analysis of *E. coli* ribosomal proteins

Ribosomal proteins are fundamental to biology because they are integral to the structure of the ribosome: Insights from NMR and X-ray crystallography about the components of the ribosome, the cellular machinery responsible for protein synthesis. Advances in various techniques have significantly enhanced our understanding of ribosome structure and function. In particular, high-resolution cryo-electron microscopy (cryo-EM) has provided detailed insights into the ribosome’s structural organization, functional mechanisms, and interactions between ribosomal proteins and RNA.^73-76^ Detailed insights into the assembly of human small ribosomal subunits have been achieved by capturing their three-dimensional structures at various stages of the assembly process.^77^ These studies have provided a comprehensive view of how these subunits are assembled. Since *E. coli* ribosomal proteins tend to degrade at 25°C after ∼24 h, it remains an unsolved task to use solution-state NMR to solve the structure of ribosome elements de-novo.^78-80^ Additionally, NMR and X-ray crystallography have been employed to reveal the flexibility and conformational dynamics of ribosomal components, further enhancing our understanding of their functional mechanisms.^81-84^ MS-based top-down proteomics (TDP) techniques have also been used to characterize *E. coli* ribosomal proteins. While these methods effectively identify different proteoforms, conventional TDP does not provide conformational states of these proteoforms.^39,85^

Up to now, there is a lack of structural information for *E. coli* ribosomal proteins in the PDB database. The structural diversity and conformational changes of the components during the assembly process present an intriguing area of study. As a demonstration, we applied the PSP-native proteomics method for analyzing *E. coli* ribosomal proteins, where protein identification and conformation determination were performed simultaneously. The ribosomal protein sample extracted by the native protein extraction workflows underwent RPLC-TDMS and SEC-nMS experiments separately. As shown in Figure 3, 54 subunit proteins were identified in the RPLC-TDMS experiment. Next, the native mass spectra of 50 corresponding subunits were referenced in the SEC-nMS experiment, allowing us to obtain the SASAs of each protein from their CSDs. Notably, ribosomal proteins were extracted as monomers rather than as part of a complete protein-RNA complex. This means that ribosomal proteins with IDRs may exhibit flexible and dynamic conformations in a near-native solvent environment, leading to the observation of multiple CSDs. Figure 3 illustrates that most ribosomal proteins display multiple CSDs and, consequently, multiple SASAs. Since the cryo-EM structure of the whole ribosome has been extensively documented, the SASA of each ribosomal protein in the ribosome complex was also listed in the scatter plot (green dots) for comparison. As shown in Figure 3, most proteins exhibit two or three SASAs, each would correspond to a minimum energy state. The four SASAs observed for L1, L13, L15, S2, S3, S4 and S12 suggest that these seven proteins are particularly dynamic and flexible. On the other hand, L29, L30, L36, and S10 each have only one SASA, suggesting that these proteins maintain in a relatively stable conformation.

**Figure 3.**
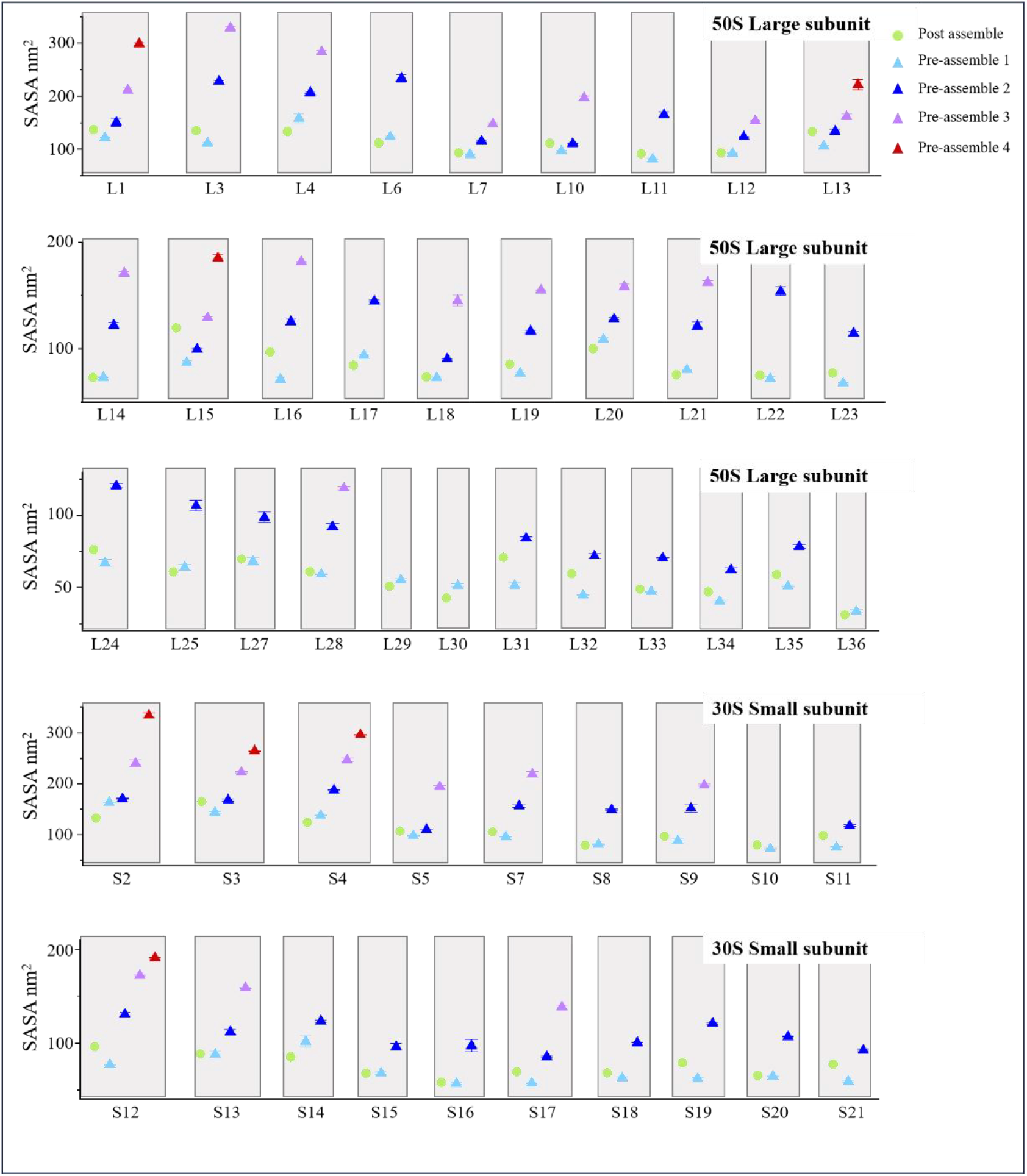
Ribosomal proteins and their SASAs measured from the RPLC-TDMS and SEC-nMS experiments. Green dots: protein SASAs when they are in the ribosome complex and extracted from the cryo-EM data. Triangles: protein SASAs obtained from their native mass spectra.

Understanding both protein conformations before and after complex formation is essential for elucidating the mechanisms of protein interactions and advancing drug development targeting these processes. As the whole ribosome is composed of ribosomal RNA and ribosomal proteins, the assembly of these molecular machines is highly complicated. It remains unclear whether the structures of individual ribosomal proteins change upon forming the ribosome complex. After acquiring protein SASAs, the conformations of each ribosomal protein were obtained and compared with their corresponding conformations in the ribosome complex. The volcano plot in Figure 4a indicates that 38% of the identified ribosomal proteins exhibit relatively minor conformational changes, most of which have fewer IDRs and more compact structures. As an example, Figure 4b compares the conformations of L25 before and after assembly, and the compact structure before assembly is almost entirely consistent with the structure after assembly.

**Figure 4.**
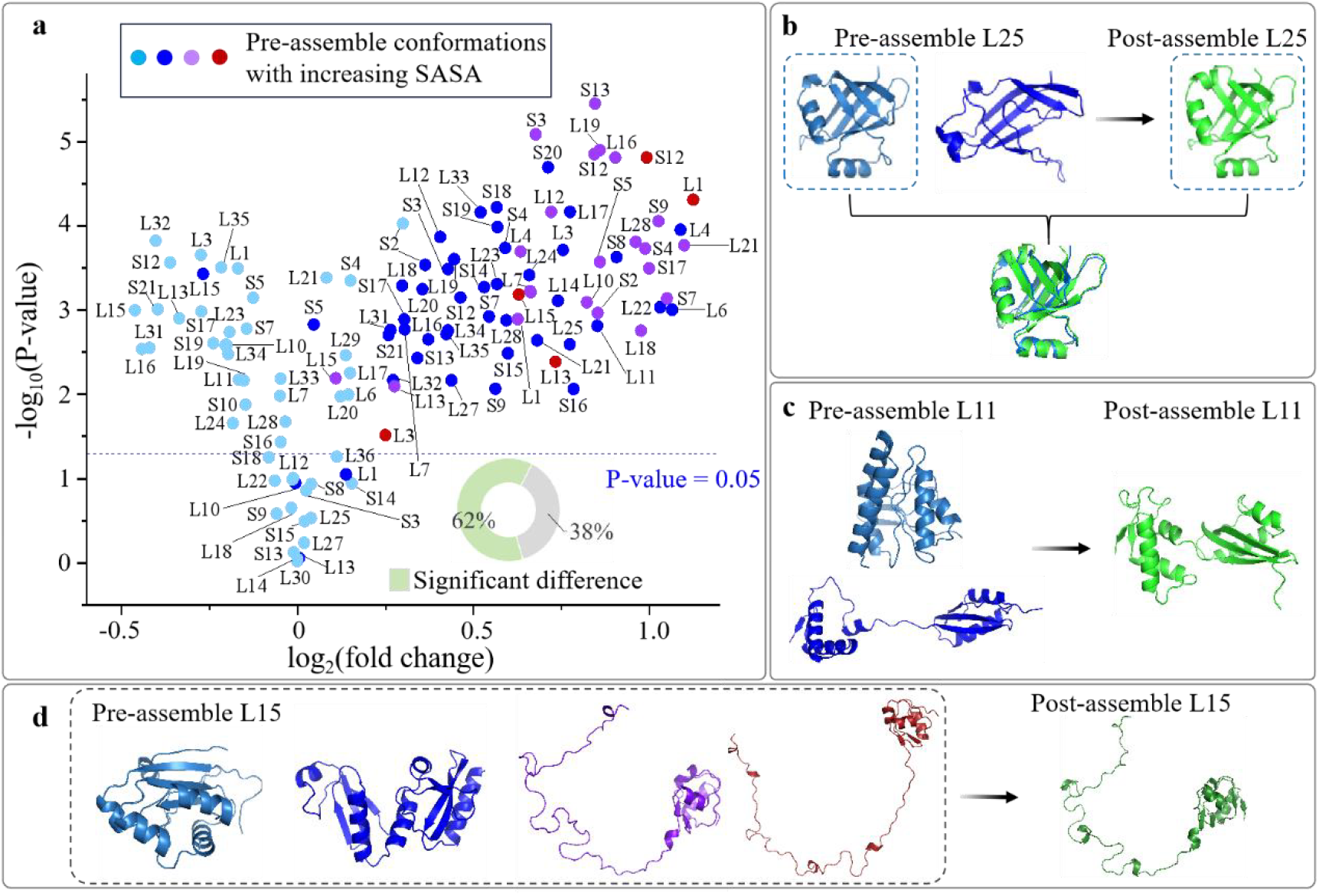
Conformation variations of ribosomal proteins pre- and post-assemble. (a) Volcano plot of ribosomal protein SASAs before and after forming complex. (b) The compact structure of L25 with no significant difference. (c) L11 undergoes a certain degree of structural transition after forming a complex. (d) The four coexisting structures of L15 in its pre-assembly state contrast with the single structure observed in its post-assembly state.

In contrast, the remaining 62% of ribosomal proteins display significant conformational variations upon integration into the ribosome assembly. These proteins often contain IDRs and can adopt multiple conformation ensembles in their monomer state. For instance, Figure 4c highlights that the compact conformations of L11 in its monomer state obviously extend upon incorporation into the ribosome complex. Figure 4d illustrates the four coexisting conformations of L15 as a monomer; however, only one conformation is observed after it is integrated into the ribosome. These findings support previous studies that IDPs can adopt various conformations in their free state, facilitating binding to targets. A dynamic equilibrium exists among these conformations, and the introduction of ligands can disrupt this balance, favoring the formation of a stable complex with the target.^86^ Consequently, protein molecules adopting other conformations shift to the favored state to support the complex formation process.

### Interactions of ribosomal proteins with a drug molecule

The protein-drug interactions (PDIs) are essential for advancing drug discovery processes. Many proteins, including ribosomal proteins, perform their biological functions by binding to other proteins or ligands. Consequently, identifying optimized molecules to block or prevent these binding events is a key objective in drug development. To achieve this, understanding the structure of the target protein before forming complex is essential. As a demonstration, kanamycin B was selected as a drug molecule, and its interactions with ribosomal proteins (before forming ribosome) were explored. In experiments, 0.1 μM kanamycin B was added to the ribosomal protein sample and incubated overnight at 4 °C. After performing nMS experiments, it is found that kanamycin B would interact with L19 protein to form the L19-kanamycin B complex. Figure 5a plots the native mass spectrum prior to the addition of kanamycin B, with the mass peaks of L19 labelled by red dots. Three Gaussian-shaped CSDs were observed, suggesting L19 could possess three conformation ensembles in its monomer state. Since these three conformations could not be resolved in SEC, it implies that individual L19 protein molecules are in a state of equilibrium, exchanging among these three conformational states on a timescale of milliseconds to microseconds.^87^ The corresponding most probable conformations of L19 are also illustrated in Figure 5a, and the compact conformation (the left structure plotted in cyan) more closely resembles that of L19 when it is within the ribosome (as shown in the Table S1).

**Figure 5.**
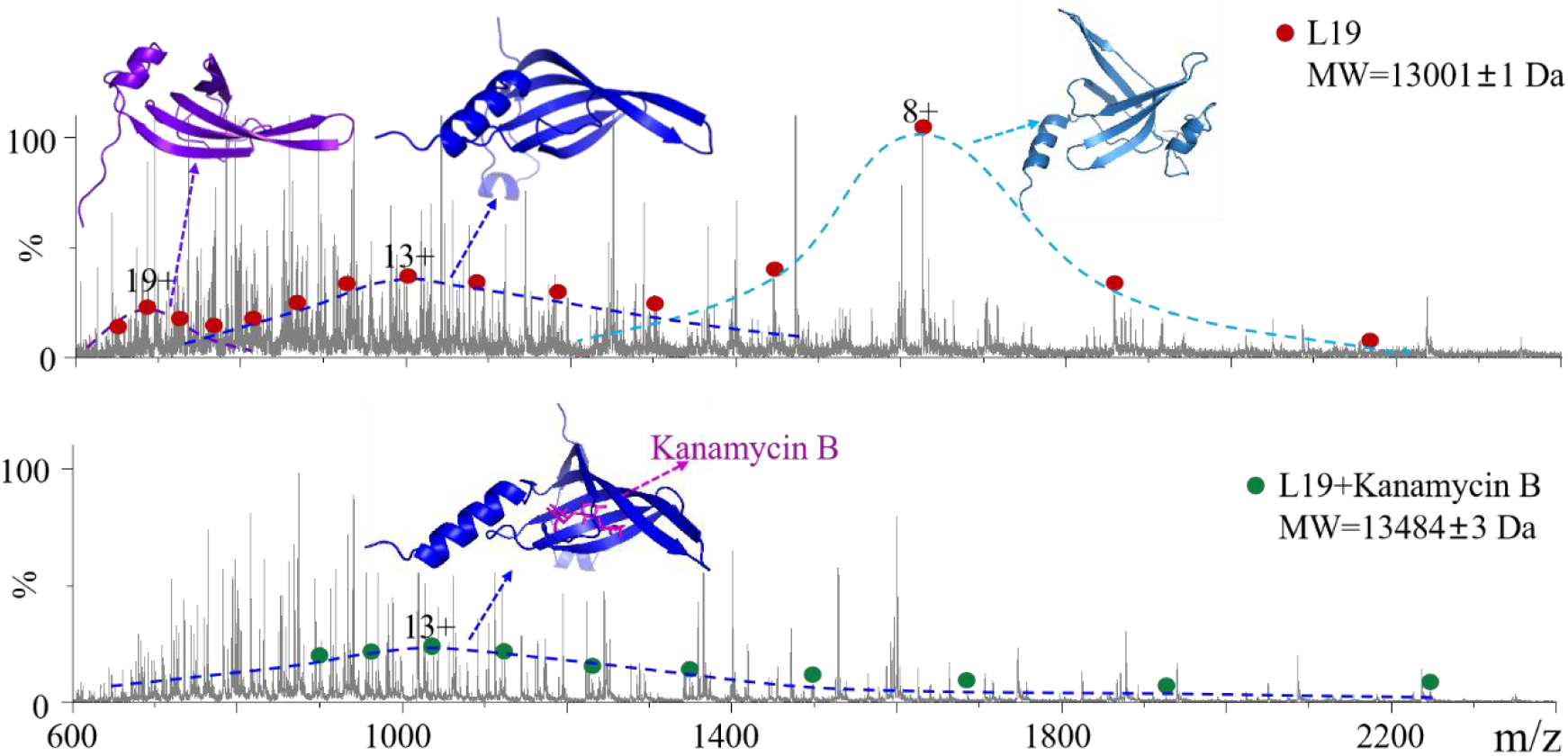
(a) Native mass spectrum of *E. coli* ribosomal proteins. Red dots annotate the mass peaks of L19 with three CSDs. Insets: conformational states of L19 corresponding to each CSD; (b) Native mass spectrum of *E. coli* ribosomal protein sample after the addition of antibiotic (kanamycin B). L19-kanamycin B complex was labeled with green dots, with its structure plotted in the inset.

After adding kanamycin B, the binding product of L19 and kanamycin B is observed in the native mass spectrum. As illustrated in Figure 5b, L19-kanamycin B complex exhibits a single CSD, and kanamycin B preferentially binds with the partially unfolded conformation of L19 (the middle structure plotted in blue). During the binding process, L19 molecules possessing the other two conformations would shift to the middle conformation to maintain equilibrium. It is important to note that the conformation of L19 that interacts with kanamycin B is not the same as the conformation of L19 within the ribosome complex. Results indicate that acquiring all possible protein conformations in the monomeric state is crucial for both the drug design process and the study of protein-ligand interactions.

## Conclusion

A comprehensive description of a proteome, encompassing both protein identities and their conformations, would greatly enhance our understanding of biological systems. We demonstrated that the AI assisted native proteomics method enables the probing of protein conformations in complex biological environments at a relatively high throughput. Protein compositions of a biological sample were acquired using the TDP method through RPLC-MS experiments, while protein conformation-related physical parameter (SASA) was measured by SEC-nMS experiments. The PSP module then utilizes both protein sequence and SASA information to output the most probable structure of each identified protein. The analysis of α-Syn and Lys show that this approach can be applied to study both IDP and globular proteins, allowing for the determination of their conformation ensembles under nMS conditions.

Notably, our application of this method to ribosomal protein analysis represents the first systematic study of ribosomal protein conformations before and after forming ribosome complex. Results revealed that most ribosomal protein with IDRs typically exhibit multiple conformation ensembles in the monomer state. Once integrated into the ribosome complex, these proteins adopt a fixed conformation. On the other hand, 19 ribosomal proteins maintain almost constant structures pre- and post-assembly. Moreover, the binding of kanamycin B with L19 suggests that the preferred protein structure a drug interacts with may not be its conformation within a protein complex.

In summary, the AI-assisted native proteomics method can be directly applied for a comprehensive characterization of the proteome in terms of both proteoforms and conformations. It facilitates the analysis of protein conformation changes before and after complex formation in complex biological samples. This technique would potentially facilitate drug developments targeting proteins.

## Method

### Materials and Chemicals

Lysozyme (Lys) was purchased from Sigma Aldrich (St. Louis, MO, USA), and α-Synuclein (α-Syn) was purchased from SinoBiological (Beijing, China). *E. coli* Ribosome (P0763S) was purchased from New England Biolabs Inc. (Ipswich, MA, USA). Ammonium acetate (CH_3_COONH_4_), glacial acetic acid (CH_3_COOH), formic acid (HCOOH), and methanol (CH_3_OH) were purchased from Dikma Co. (Beijing, China). Kanamycin B was purchased from Targetmol Co. (Boston, Massachusetts, USA). Deionized water was obtained from Wahaha Co. (Hangzhou, China). An ACQUITY UPLC Protein BEH SEC Column (200 Å, 1.7 μm, 2.1 mm × 150 mm) was used to separate ribosomal proteins in the SEC-nMS experiment. An ACQUITY UPLC BEH300 C4 column (1.7 μm, 2.1 mm × 50 mm) was used in the RPLC-TDMS experiment. Molecular weight cutoff (MWCO) filters at 3 kDa were purchased from Millipore (Billerica, MA, USA).

### Sample Preparement

Lysozyme and α-Synuclein were dissolved in 100 mM ammonium acetate at a concentration of 0.7 μM. 200 μL glacial acetic acid was added to the *E. coli* ribosome vial. The ribosome sample was incubated at 4 °C overnight, and then centrifuged at 12,000 rpm for 10 minutes to pellet the rRNA. The ribosomal sediment was removed, and the ribosomal proteins were collected in the supernatant. Ultrafiltration was performed using 3 kDa molecular weight cutoff (MWCO) filters three times to exchange the buffer with the 25 mM ammonium acetate solution. 10 μM kanamycin B was added in ribosomal protein solution, and the mixture of protein and drug was incubated overnight at 4 °C.

### SEC-nMS experiment

A SEC-nMS experiment was conducted using the Acquity ultra-performance liquid chromatography (UPLC) I-Class system coupled with a Xevo G2-XS QTOF mass spectrometer (Waters Corporation, Wilmslow, UK). A neutral mobile phase and gentle MS parameters were used to maintain the native structure of proteins. Specifically, a 25 mM ammonium acetate solution was used as the mobile phase. The flow rate of the mobile phase was set to 0.1 mL/min. As for the MS parameters, the capillary voltage was set at 1.5 kV, and the source temperature was set to 70 °C. The desolvation temperature was set to 400 °C, and the desolvation gas flow was set to 600 L/h. The injection volume of the ribosomal protein sample was 10 μL, whereas the injection volumes of α-Synuclein and lysozyme were 2 μL each.

### RPLC-TDMS experiment

For the analysis of the ribosomal protein sample, an ACQUITY UPLC BEH300 C4 column (1.7 μm, 2.1 mm ×50 mm) was used. Mobile phase A (MPA) was 0.1% formic acid in water, and mobile phase B (MPB) was 0.1% formic acid in methanol. The gradient of MPA was set to 95%, 58%, 5%, and 95% at 0, 65, 70, and 80 minutes, respectively. The entire run time was 90 minutes. The flow rate of the mobile phase was 0.2 mL/min. The injection volume for all samples was 2 μL. The MS was operated in positive and MSe (data-independent acquisition) mode. Additionally, a ramp high-energy collision dissociation (HCD) from 20-70 V was used to fragment ribosomal protein ions.

As there is currently no suitable software available for processing data collected using full-window DIA mode in top-down proteomics, the RPLC-TDMS data of ribosomal proteins were processed utilizing a self-developed data processing procedure.^63^ Initially, the raw data file was converted to the .mzML format using MSConvert software. Subsequently, the .mzML files underwent preprocessing through a deep learning-based model designed to eliminate noise signals. Following this step, the denoised mass spectrum was analyzed using Unidec, FlashDeconv, and TopPIC software for deconvolution. By evaluating and integrating results from these three software tools, an ion pairing strategy was employed to elucidate relationships between precursors and fragments. At this stage, a pseudo-msalign file was generated. Lastly, the msalign files were analyzed by TopPIC for database searching. The mass error tolerance was set to 30 ppm, the maximum number of mass shifts was set to 2 Da, and the allowed mass shift was set to 500 Da.

### PSP AI module

This research employed a STR2STR (Structure-to-Structure Translation) framework, a score-based method designed for zero-shot protein conformation sampling.^61^ This methodology is trained exclusively on general protein structures and facilitates efficient conformation sampling of “dark proteins” without dependence on simulation data specific to the target.^62,88^

The fundamental concept behind the STR2STR approach involves achieving protein conformation sampling through a denoising diffusion-based forward-backward procedure. Specifically, the method introduces stochastic perturbations to the input structure in the forward phase. Subsequently, it employs a backward phase guided by a scoring function to progressively remove the introduced noise. This process is inspired by simulated annealing and allows for the effective generation of a diverse ensemble of protein backbone conformations while preserving global rotational and translational invariances.

To translate the protein backbone structures generated by STR2STR into full-atom models, the CG2ALL (Coarse-Grained to All-Atoms) method was applied. CG2ALL represents an efficient and precise approach for refining coarse-grained protein backbone structures into detailed all-atom representations.

## Conflicts of interest

There are no conflicts to declare.

### Acknowledgements

This work was supported by NNSFC (22374009).

## Contributions

W.X. contributed to conceptualization, supervision, management, manuscript reviewing and editing. W.Z. contribute to the entire experiments, analyzed the SEC-nMS data and drafted the manuscript. C.S. analyzed the RPLC-TDMS data. Z.X contributed to employ STR2STR (Structure-to-Structure Translation) framework to generate the structure ensemble.

## Supporting Information

The Supporting Information is available free of

