## Supplemental Table S1 for "AI Assisted Native Proteomics: Delineating Ribosomal Protein Conformations Pre- and Post-Assembly"

### Table S1 The structures of each ribosomal protein pre- and post- assembly

| Description | Uniprot Accession | Post-assemble | Pre-assemble 1 | Pre-assemble 2 | Pre-assemble 3 | Pre-assemble 4 |
|---|---|---|---|---|---|---|
| 50S ribosomal protein L1 | A0A140NFM7 | 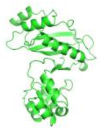 | 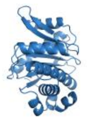 | 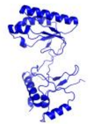 | 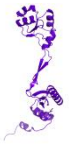 | * |
| 50S ribosomal protein L3 | A0A140N3G7 | 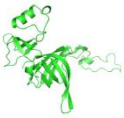 | 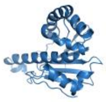 | 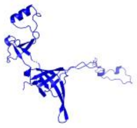 | * | |
| 50S ribosomal protein L4 | A0A140N5K8 | 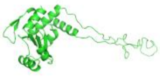 | 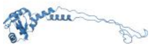 | 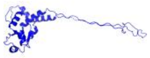 | * | |
| 50S ribosomal protein L6 | A0A140N2T1 | 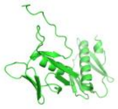 | 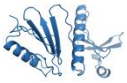 | 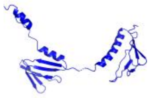 | | |
| 50S ribosomal protein L7 | A0A140SS63 | 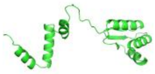 | 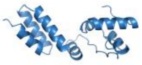 | 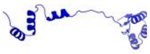 | * | |
| 50S ribosomal protein L10 | A0A140NDB6 | 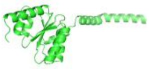 | 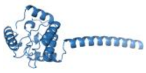 | 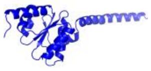 | 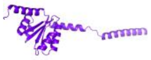 | |
| 50S ribosomal protein L11 | A0A140NF32 | 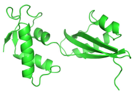 | 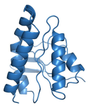 | 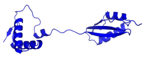 | | |
| 50S ribosomal protein L12 | A0A140SS63 | 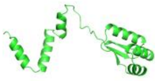 | 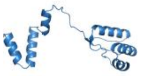 | 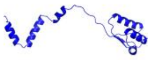 | * | |
| 50S ribosomal protein L13 | A0A140N598 | 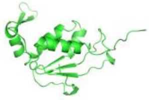 | 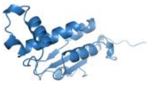 | 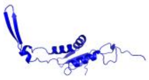 | * | * |
| 50S ribosomal protein L14 | A0A140N3H4 | 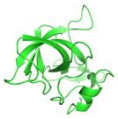 | 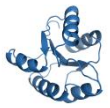 | 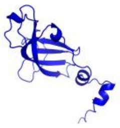 | * | |

| Description | Uniprot Accession | Post-assemble | Pre-assemble 1 | Pre-assemble 2 | Pre-assemble 3 | Pre-assemble 4 |
|---|---|---|---|---|---|---|
| 50S ribosomal protein L15 | A0A140N711 | 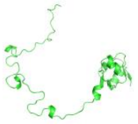 | 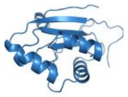 | 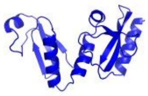 | 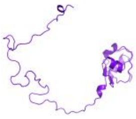 | 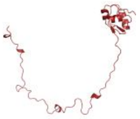 |
| 50S ribosomal protein L16 | A0A140N6Z2 | 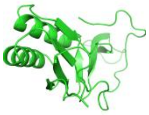 | 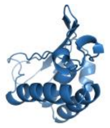 | 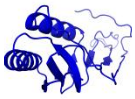 | * | |
| 50S ribosomal protein L17 | A0A140N4M0 | 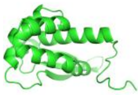 | 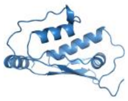 | 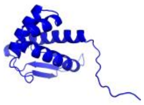 | | |
| 50S ribosomal protein L18 | A0A140N4L0 | 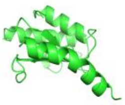 | 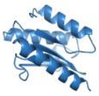 | 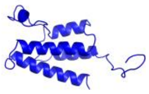 | * | |
| 50S ribosomal protein L19 | A0A140N6T7 | 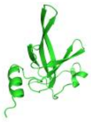 | 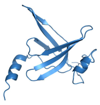 | 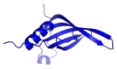 | 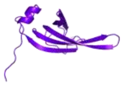 | |
| 50S ribosomal protein L20 | A0A140NBS1 | 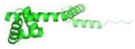 | 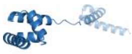 | 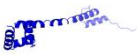 | * | |
| 50S ribosomal protein L21 | A0A140N5D7 | 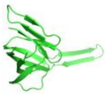 | 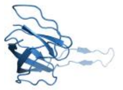 | 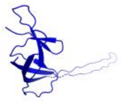 | * | |
| 50S ribosomal protein L22 | A0A140N2S3 | 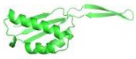 | 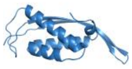 | 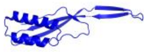 | | |
| 50S ribosomal protein L23 | C6EGE8 | 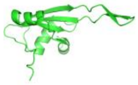 | 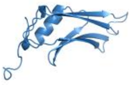 | 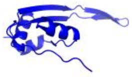 | | |
| 50S ribosomal protein L24 | A0A140N5L7 | 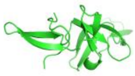 | 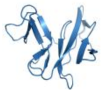 | 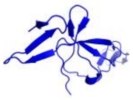 | | |

| Description | Uniprot Accession | Post-assemble | Pre-assemble 1 | Pre-assemble 2 | Pre-assemble 3 | Pre-assemble 4 |
|---|---|---|---|---|---|---|
| 50S ribosomal protein L25 | A0A140N846 | 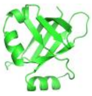 | 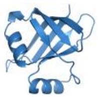 | 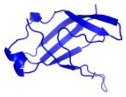 | | |
| 50S ribosomal protein L27 | A0A140N340 | 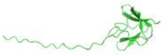 | 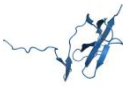 | 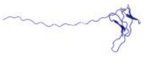 | | |
| 50S ribosomal protein L28 | A0A140N3L9 | 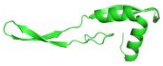 | 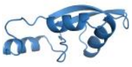 | 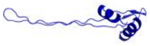 | * | |
| 50S ribosomal protein L29 | A0A140N7F5 | 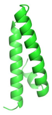 | 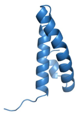 | | | |
| 50S ribosomal protein L30 | A0A140N7G4 | 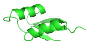 | 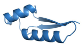 | | | |
| 50S ribosomal protein L31 | A0A140SS71 | 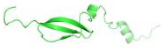 | 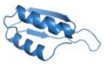 | 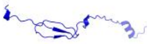 | | |
| 50S ribosomal protein L32 | A0A140NAZ2 | 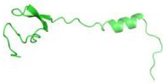 | 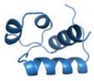 | 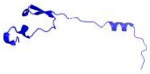 | | |
| 50S ribosomal protein L33 | A0A140N5Z7 | 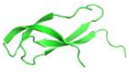 | 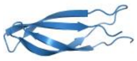 | 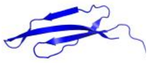 | | |
| 50S ribosomal protein L34 | A0A140NHV0 | 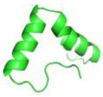 | 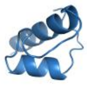 | 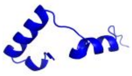 | | |
| 50S ribosomal protein L35 | A0A140N9E3 | 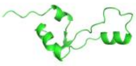 | 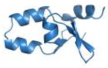 | 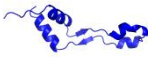 | | |

| Description | Uniprot Accession | Post-assemble | Pre-assemble 1 | Pre-assemble 2 | Pre-assemble 3 | Pre-assemble 4 |
|---|---|---|---|---|---|---|
| 50S ribosomal protein L36 | A0A140N5M6 | 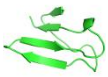 | 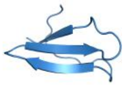 | | | |
| 30S ribosomal protein S2 | A0A140NFK2 | 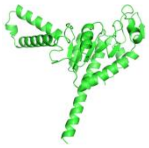 | 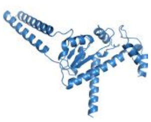 | 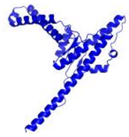 | 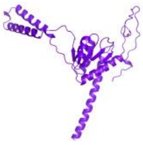 | * |
| 30S ribosomal protein S3 | A0A140N4K1 | 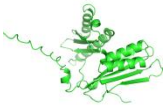 | 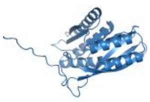 | 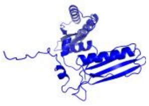 | 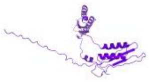 | * |
| 30S ribosomal protein S4 | A0A140N548 | 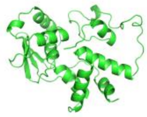 | 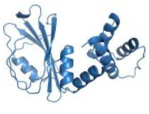 | 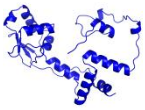 | * | * |
| 30S ribosomal protein S5 | A0A140N6Z9 | 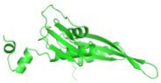 | 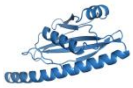 | 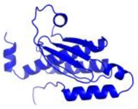 | 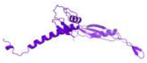 | |
| 30S ribosomal protein S7 | A0A140N6W8 | 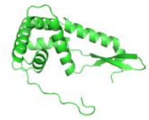 | 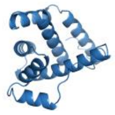 | 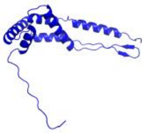 | * | |
| 30S ribosomal protein S8 | A0A140N537 | 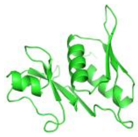 | 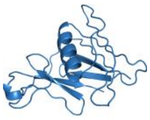 | 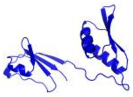 | | |
| 30S ribosomal protein S9 | A0A140N2Z9 | 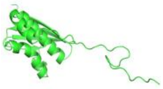 | 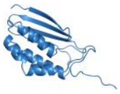 | 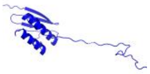 | * | |
| 30S ribosomal protein S10 | A0A140N6Y5 | 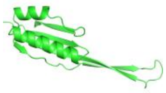 | 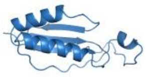 | | | |
| 30S ribosomal protein S11 | A0A140N7L9 | 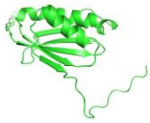 | 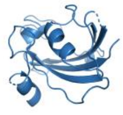 | 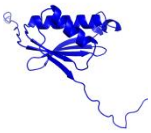 | | |

| Description | Uniprot Accession | Post-assemble | Pre-assemble 1 | Pre-assemble 2 | Pre-assemble 3 | Pre-assemble 4 |
|---|---|---|---|---|---|---|
| 30S ribosomal protein S12 | A0A140N4H3 | 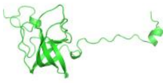 | 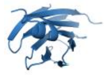 | 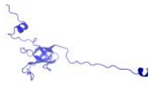 | * | * |
| 30S ribosomal protein S13 | A0A140N5B4 | 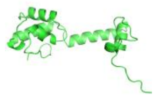 | 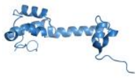 | 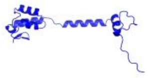 | * | |
| 30S ribosomal protein S14 | A0A140N7K8 | 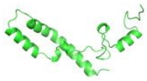 | 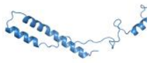 | 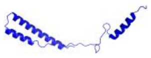 | | |
| 30S ribosomal protein S15 | A0A140N811 | 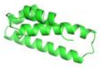 | 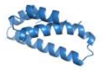 | 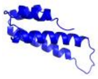 | | |
| 30S ribosomal protein S16 | A0A140N7D8 | 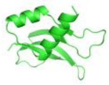 | 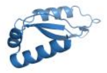 | 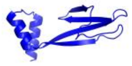 | | |
| 30S ribosomal protein S17 | A0A140N6Z6 | 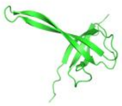 | 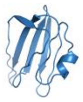 | 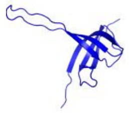 | * | |
| 30S ribosomal protein S18 | A0A140NGH1 | 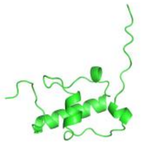 | 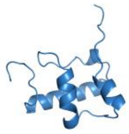 | 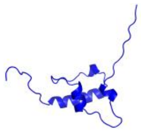 | | |
| 30S ribosomal protein S19 | A0A140N528 | 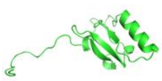 | 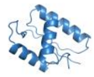 | 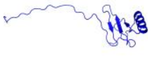 | | |
| 30S ribosomal protein S20 | A0A140NFU3 | 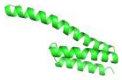 | 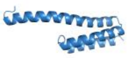 | 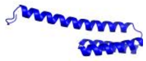 | | |
| 30S ribosomal protein S21 | A0A140N8B4 | 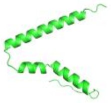 | 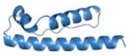 | 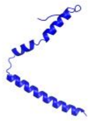 | | |

* The structure associated with the SASA value was not found within the STR2STR system.
